# Development of a modular automated system for maintenance and differentiation of adherent human pluripotent stem cells

**DOI:** 10.1101/079335

**Authors:** Duncan E. Crombie, Maciej Daniszewski, Helena H. Liang, Tejal Kulkarni, Fan Li, Grace E. Lidgerwood, Alison Conquest, Damian Hernández, Sandy S. Hung, Katherine P. Gill, Elisabeth De Smit, Lisa S. Kearns, Linda Clarke, Valentin M. Sluch, Xitiz Chamling, Donald J. Zack, Raymond C.B. Wong, Alex W. Hewitt, Alice Pébay

**Author notes:** Co-first authors (Duncan E. Crombie, Maciej Daniszewski, Helena H. Liang). Co-senior authors (Alex W. Hewitt, Alice Pébay). Corresponding author: Alice Pébay, PhD, Address: CERA, University of Melbourne, 32 Gisborne Street, East Melbourne, VIC 3002, Australia. Address for correspondence: Alice Pébay. University of Melbourne, 32 Gisborne Street, East Melbourne, VIC 3002 Australia.

## Abstract

Patient-specific induced pluripotent stem cells (iPSCs) have tremendous potential for development of regenerative medicine, disease modelling and drug discovery. However, the processes of reprogramming, maintenance and differentiation are labour intensive and subject to inter-technician variability. To address these issues, we established and optimised protocols to allow for the automated maintenance of reprogrammed somatic cells into iPSCs to enable the large-scale culture and passaging of human pluripotent stem cells (PSCs) using a customized TECAN Freedom EVO. Generation of iPSCs was performed offline by nucleofection followed by selection of TRA-1-60 positive cells using a Miltenyi MultiMACS24 Separator. Pluripotency markers were assessed to confirm pluripotency of the generated iPSCs. Passaging was performed using an enzyme-free dissociation method. Proof of concept of differentiation was obtained by differentiating human PSCs into cells of the retinal lineage. Key advantages of this automated approach are the ability to increase sample size, reduce variability during reprogramming or differentiation, and enable medium to high-throughput analysis of human PSCs and derivatives. These techniques will become increasingly important with the emergence of clinical trials using stem cells.

## Introduction

Advances in technology have enabled the reprogramming of adult somatic cells into human induced pluripotent stem cells (iPSCs) (Park et al. 2008; Takahashi et al. 2007; Yu et al. 2007), which can be subsequently differentiated into cells of interest, providing a potentially inexhaustible supply of cells with disease-specific genotypes and phenotypes. A main bottleneck in using iPSCs for disease modeling lies in the vast amount of time and manpower necessary to maintain cells in culture, making large-scale population studies almost unfeasible within the constraints of an average sized laboratory. The use of robotics could enable this type of research. A key advantage of an automated approach is the ability to increase sample size and reduce variability during the reprogramming, maintenance and differentiation of iPSCs. Current applications of automated technology for stem cell research are still limited. Among the very few systems reported for automation of stem cell culture and differentiation are the Automation Partnership Biosystems (TAP Biosystems), which has already been used to maintain human mesenchymal stem cells (Thomas et al. 2008a) as well as human bone marrow-derived cells (Thomas et al. 2008b); and the AutoCulture^®^ system (Kawasaki Heavy Industries), which has been used to maintain human cardiac stem cells (Kami et al. 2013). Additionally, prototypes have been developed to change medium for cell culture of embryonic stem cells (Reichen et al. 2013). The automated platform TECAN Freedom EVO (TECAN) has been adapted to maintain mouse embryonic stem cells and differentiate them towards a neuronal lineage (Hussain et al. 2013). Similarly, a few groups have reported on the reprogramming, maintenance and differentiation of human PSCs on automated platforms. The New York Stem Cell Foundation uses a self-designed robotic platform comprised of three platforms using STAR liquid handling systems (Hamilton Robotics) to carry-out the maintenance and passaging of human iPSCs (Paull et al. 2015). Another automated platform was described for the maintenance of human iPSCs, using a robotic arm (MELFA, RV-4FC-D, Mitsubishi) for liquid handling (Konagaya et al. 2015). A smaller liquid handler was recently described for the maintenance and passaging of human PSCs, based on a self-contained Gilson’s pipette Max liquid handler, that can process up to 96 well plates (Conway et al. 2015). However, this system requires additional offline steps as there is no associated incubator and many steps require human contribution.

Herein we describe a customized TECAN Freedom EVO platform for the maintenance of human fibroblasts undergoing reprogramming to iPSCs, as well as maintenance and passaging of undifferentiated colonies of PSCs. We also report the feasibility of using this platform for long term differentiation of cells, demonstrated with guided differentiation of human PSCs to retinal cells, including retinal ganglion cells (RGCs) and retinal pigment epithelium (RPE) cells. We describe our maintenance protocols employed and the adaptations required for using this automated platform. This system allows for large sample sized research, reduced variability and allows for future high-throughput analysis of the transcriptome and metabolome of progeny cells derived from patient iPSCs (Suppl. Fig. 1).

## Materials and Methods

### Ethics

All experimental work performed in this study was approved by the Human Research Ethics committees of the Royal Victorian Eye and Ear Hospital (11/1031H, 13/1151H-004) and University of Melbourne (0605017, 0829937) with the requirements of the National Health & Medical Research Council of Australia (NHMRC) and conformed with the Declarations of Helsinki (McCaughey et al. 2016).

### Platform material

The TECAN system is a liquid handling platform that requires a tissue culture plate format. All cells were cultured and handled in a 6-well plate format. Tips used were 5 mL disposable conductive sterile tips with filter for LiHa arm (TECAN). Cell culture medium was aliquoted into 50 mL falcon tubes and placed into specific carriers in the TECAN platform.

### Fibroblast Culture

Human fibroblasts were cultured in DMEM with high glucose, 10% fetal bovine serum (FBS), L-glutamine, 100 U/mL Penicillin and 100 µg/mL streptomycin (all from Life Technologies). All cell lines were confirmed to be mycoplasma-free using the MycoAlert mycoplasma detection kit (Lonza) according to the manufacturer’s instructions.

### iPSC generation

iPSCs were generated using human skin fibroblasts obtained from subjects over the age of 18 years by episomal method as described previously (Hung et al. 2016). Briefly, reprogramming was performed on passage 8-10 fibroblasts by nucleofection (Lonza Amaxa Nucleofector) with episomal vectors expressing OCT4, SOX2, KLF4, L-MYC, LIN28 and shRNA against p53 (Okita et al. 2011) in feeder- and serum-free conditions using TeSR-E7 medium (Stem Cell Technologies). The reprogrammed cells were maintained on the automated platform using TeSR-E7 medium, with medium change every day.

### PSC Selection, maintenance and passaging of iPSCs

Pluripotent cells were selected using a MultiMACS (Miltenyi) by TRA-1-60 sorting using anti-human TRA-1-60 Microbeads in combination with MultiMACS Cell24 Columns (Miltenyi). Briefly, reprogrammed cells from 1 well of a 6-well multiwell plate were incubated with TrypLE (5-10 mins, 37°C), cells were collected and gently triturated in TeSR-E8 medium supplemented with Y27632 (10 µM). The cell suspension was filtered through a pre-separation filter (30 µm, Miltenyi), centrifuged and resuspended in 80 µL ice-cold TeSR-E8 medium containing Y27632 (10 µM), and incubated with 20 µL anti-TRA-1-60 beads (5 mins, 4°C). Volume was adjusted to 1 mL in TeSR-E8 medium containing Y27632 (10 µM) and each suspension containing magnetically labelled cells was loaded onto a MultiMACS column. Columns were washed twice with TeSR-E8 medium containing Y27632, then eluted with 1mL TeSR-E8 medium containing Y27632. Cell number was determined and cells were then plated into 1 well of a 6-well plate coated with vitronectin XF (40 µL/well in 2 mL cell adhere dilution buffer Stem Cell Technologies), then placed back into the online incubator. Subsequent culturing was performed on the automated platform using TeSR-E8 (Stem Cell Technologies), changing medium every two days. Passaging of the iPSC line CERA 007 (Hernandez et al. 2016) was performed on the automated platform using ReLeSR^TM^ (Stem Cell Technologies) onto vitronectin XF plated wells.

### Retinal cell differentiation

Retinal differentiation of BRN3B-mCherry A81-H7 hESCs (Sluch et al. 2015) was performed via an adapted protocol originally described by (Lamba et al. 2006) using DMEM F12 with glutaMAX (Life Technologies), 10% Knockout Serum Replacement (Life Technologies), IGF1 (10 ng/mL, Peprotech), Dkk1 (10 ng/mL, Peprotech), Noggin (10 ng/mL, R&D Systems), bFGF (5 ng/mL), B27 and N2 (both 1x, Life Technologies) as described in (Gill et al. 2016), changing medium every second day. We adapted the protocol to automation by starting with a monolayer of PSCs plated on vitronectin XF in place of embryoid body formation. Cells were assessed at day 24 and no further enrichment was performed (Gill et al. 2016). Successful differentiation into RGCs was determined by appearance of mCherry positive cells, which is indicative of BRN3B expression. Differentiation of H9 hESCs (Wicell, USA) into RPE cells was performed in feeder-free conditions as described in (Lidgerwood et al. 2016) using vitronectin XF and RPEM medium (*α*-MEM, 0.1 mM Non Essential Amino Acids, 0.1 mM N2, 1 % L-Glutamine–Penicillin–Streptomycin solution, 250 µg/mL Taurine, 20 ng/mL Hydrocortisone, 13 pg/mL Triiodothyronine (all from Sigma-Aldrich), 25 mM HEPES), supplemented with 5% FBS, IGF1 (10 ng/mL), Dkk1 (10 ng/mL), Noggin (10 ng/mL), bFGF (5 ng/mL), B27 and N2 (both 1x), changing medium every two days. Cells were assessed at day 35. Successful differentiation into RPE cells was determined by cobblestone morphology and pigmentation, as well as PMEL expression by immunocytochemistry.

### Immunocytochemistry

Immunocytochemistry was performed using OCT3/4 (C-10, Santa Cruz), TRA-1-60 (Abcam) and PMEL (Abcam). Cells were then immunostained with isotype-specific secondary antibodies (Alexa-Fluor, Life Technologies). Nuclei were counterstained using DAPI (Sigma-Aldrich).

## Results

### Description of the automated platform

Our modular platform is comprised of a TECAN Freedom EVO 200 - that includes a class 2 biosafety cabinet, a robotic liquid handling arm with 8 independent channels (8 channels LiHa) and a robotic manipulator arm (RoMa with eccentric fingers) - in conjunction with a Liconic STX110 automated incubator mounted behind the Freedom EVO, and a carousel LPX220 for dispenser tips (DiTis) on the right side of the working platform (Fig. 1). A 5 L glass bottle and peristaltic pumps are also present; however, these were not used for the routine maintenance described here (Fig. 1). The overall system dimensions are 3.4m (L) × 1.8 m (W) × 2.7 m (H). To the left side, some space was reserved for a computer, keyboard, monitor and operator (Fig. 1). A MultiMACS24 Separator and a MACSquant flow cytometer were accessed offline. As illustrated in Figure 1, on the workbench, from left to right, are a cabinet with medium bottle and pump (1) which can deliver medium to a LiHa medium refill trough contained within a Torrey Pines heater; 3 carriers for up to 24 × 50 mL falcon tubes (2), a LiHa wash station (3), a carrier for 5 mL DiTis (4), a large waste for DiTi Boxes and used tips (5), a carrier for tip boxes (6), a tilting carrier (7), a microplate carrier (8), the transfer station (9) to the Liconic incubator (12), a hotel for tissue culture lids (10) and a carousel (11). Autoclaved water bottles for the liquid handling system were stored under the bench (13). The LiHa arm was used solely with 5 mL syringes: 8 × 5mL syringes for DiTis, though are capable of handling smaller 1 mL tips (which were not used for our protocols). The Liconic LPX220 carousel was used for the storage of tip boxes. It contains 1 rotary plate with 10 interchangeable cassettes and an internal robotic handler, as well as a transfer station to place tip boxes onto the worktable. The Liconic STX110 incubator comprises of an internal robotic handler to access 5 independent stackers that can store up to 85 culture plates (17 per stacker), an internal barcode scanner as well as a transfer station to bring culture plates onto the worktable. The incubator has a controlled environment, which was set to 37℃ and 5% CO_2_. The TECAN EVO runs with two independent software packages: the freedom EVOware Plus pipetting software (EVOware) and the Workflow Planning Tool. These are used to direct pipetting and device commands (EVOware) and to plan and execute each line of the worksation’s workflow (the Workflow Planning Tool). Each protocol used on the platform was entered as an independent template, as shown in Table 1.

**Figure 1.**
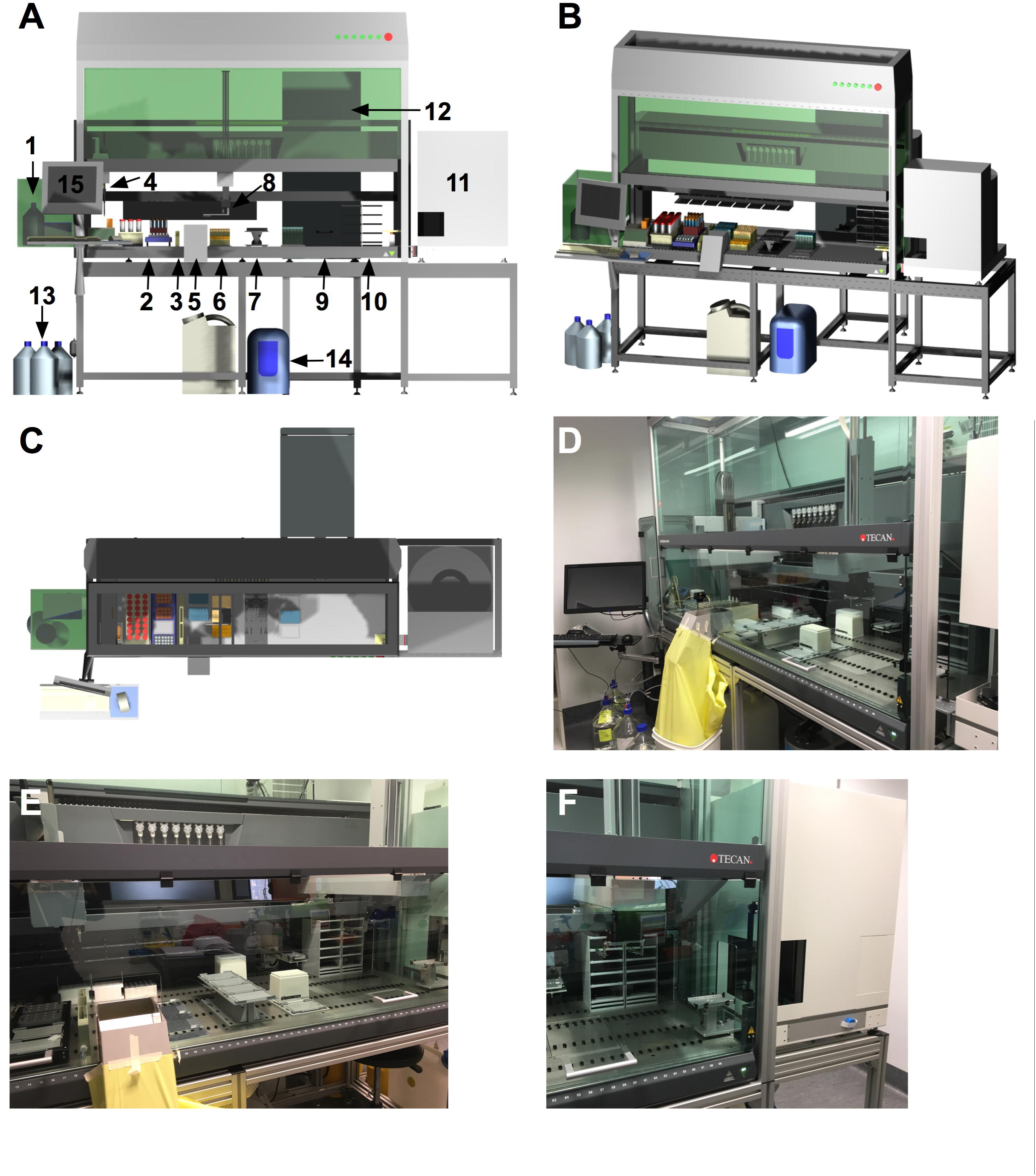
The automated platform. **(A-C)** Computer Aided Design (CAD) images of the platform from front **(A)**, angled **(B)** and top **(C)**. **(D-F)** Images of the platform. (1) cabinet with medium bottle and pump; (2) carriers for tubes; (3) LiHa wash station; (4) carrier for 5 mL DiTis; (5) waste; (6) carrier for tip boxes; (7) tilting carrier; (8) microplate carrier; (9) transfer station to the Liconic incubator; (10) hotel; (11) carousel, (12) incubator; (13) autoclaved water bottles; (14): cooling system; (15): computer.

**Table 1.**
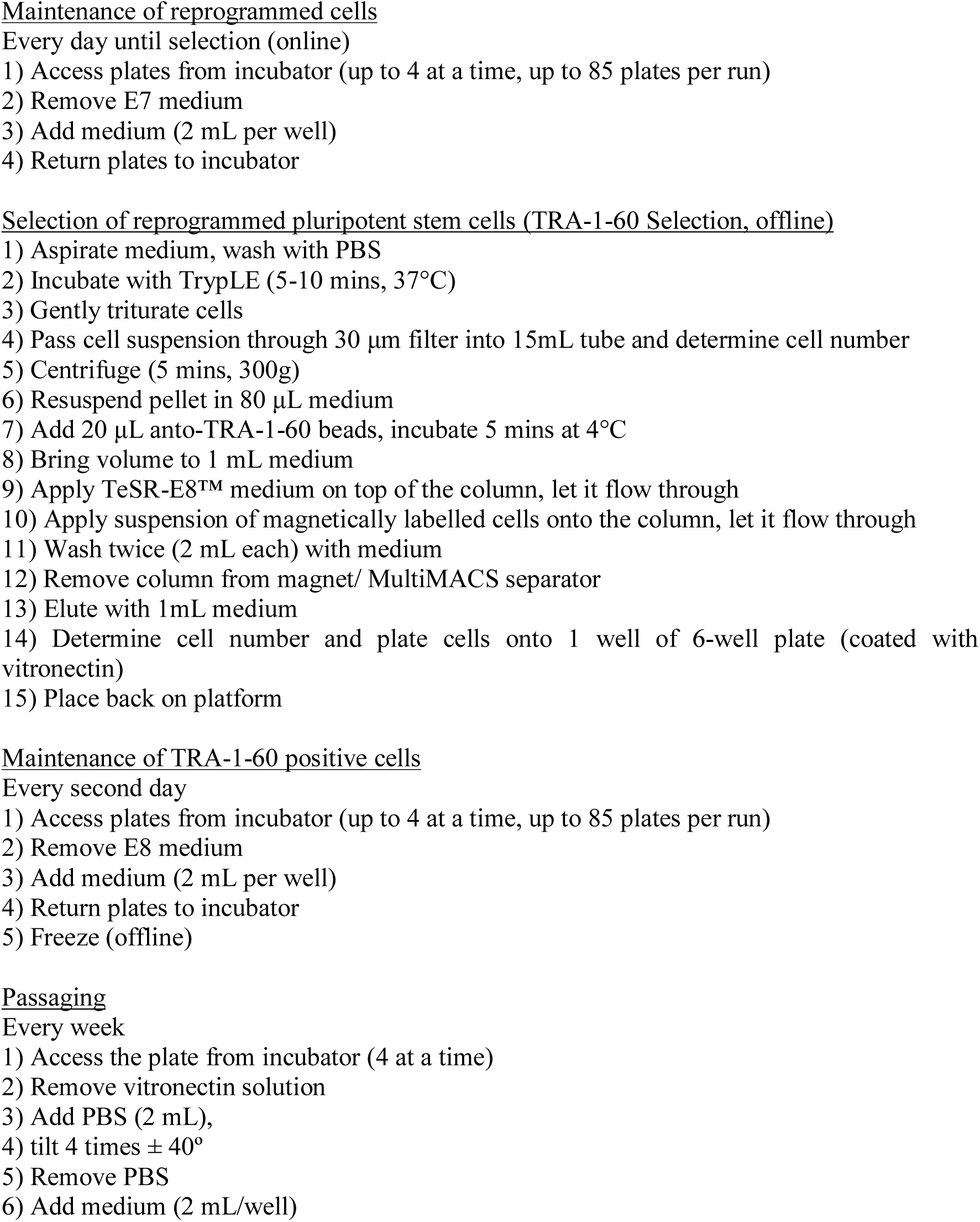
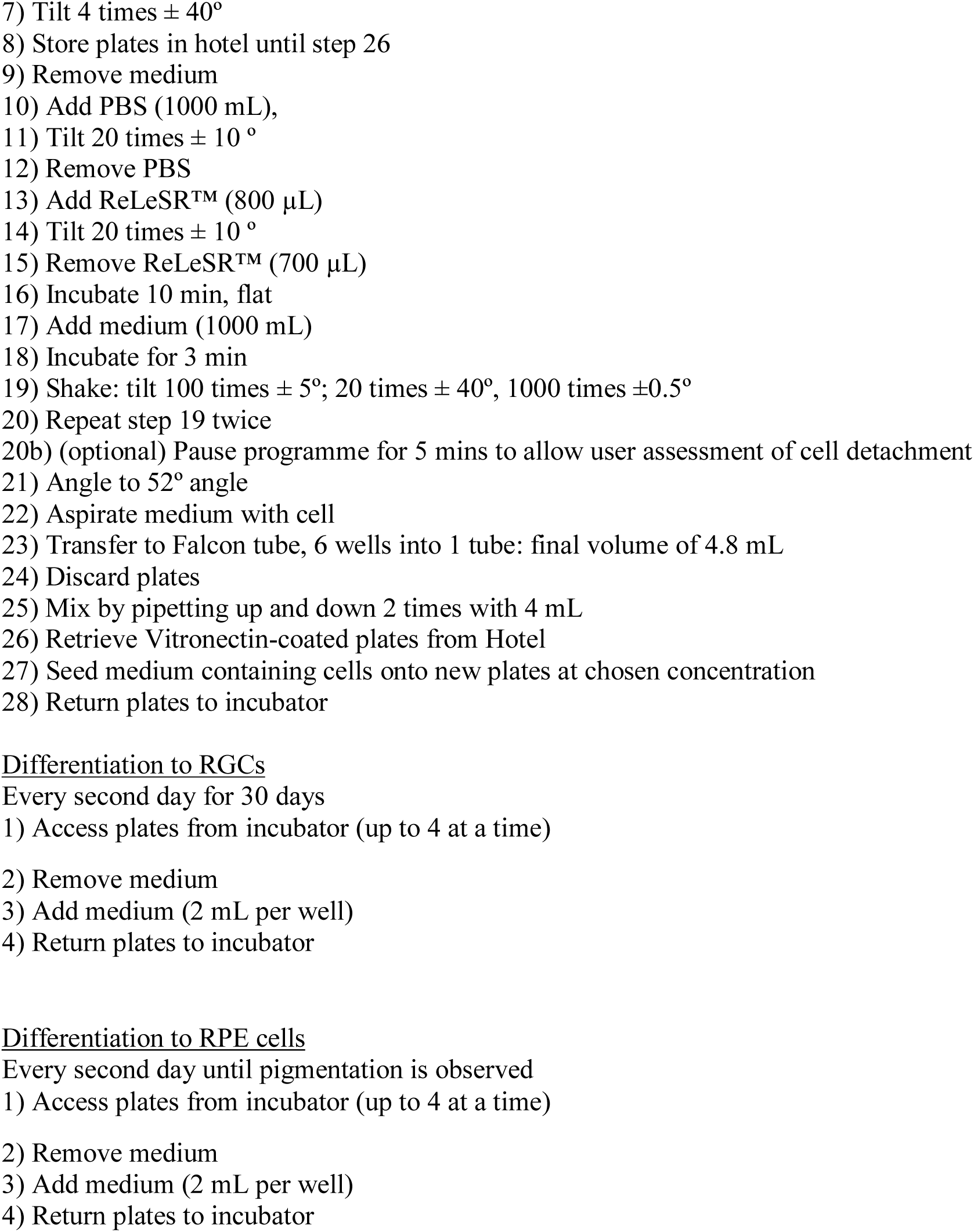
Workflow of the automated platform. Each protocol (maintenance od undifferentiated cells, passaging, RGC differentiation, RPE cell differentiation) was programmed as an independent template.

### Generation of iPSC lines

We manually reprogrammed skin fibroblasts to iPSCs using episomal vectors in feeder- and serum-free conditions in TeSR-E7 medium. Nucleofection of fibroblasts was performed in a 6-well plate format. Following nucleofection, cells were placed into the online incubator. Medium was changed every day using the automated platform. To identify and isolate iPSCs, we utilized the marker TRA-1-60 which was previously shown to be a marker of fully reprogrammed iPSCs (Chan et al. 2009). Instead of picking clonal-derived iPSCs, we performed bulk selection of polyclonal iPSCs as these were shown to be indistinguishable from clonal-derived iPSCs. Notably, the bulk generation of polyclonal iPSCs has been shown to be as effective in the generation of fully reprogrammed lines as manual selections of clones (Willmann et al. 2013). At approximately day 30, iPSCs were isolated by MACS based on TRA-1-60 positive selection using a MultiMACS24 Separator and maintained in feeder-free culture on vitronectin in TeSR-E8 medium. Pluripotency of all derived iPSC lines were confirmed by immunocytochemistry for OCT-4 and TRA-1-60 expression (Fig. 2A-F).

**Figure 2.**
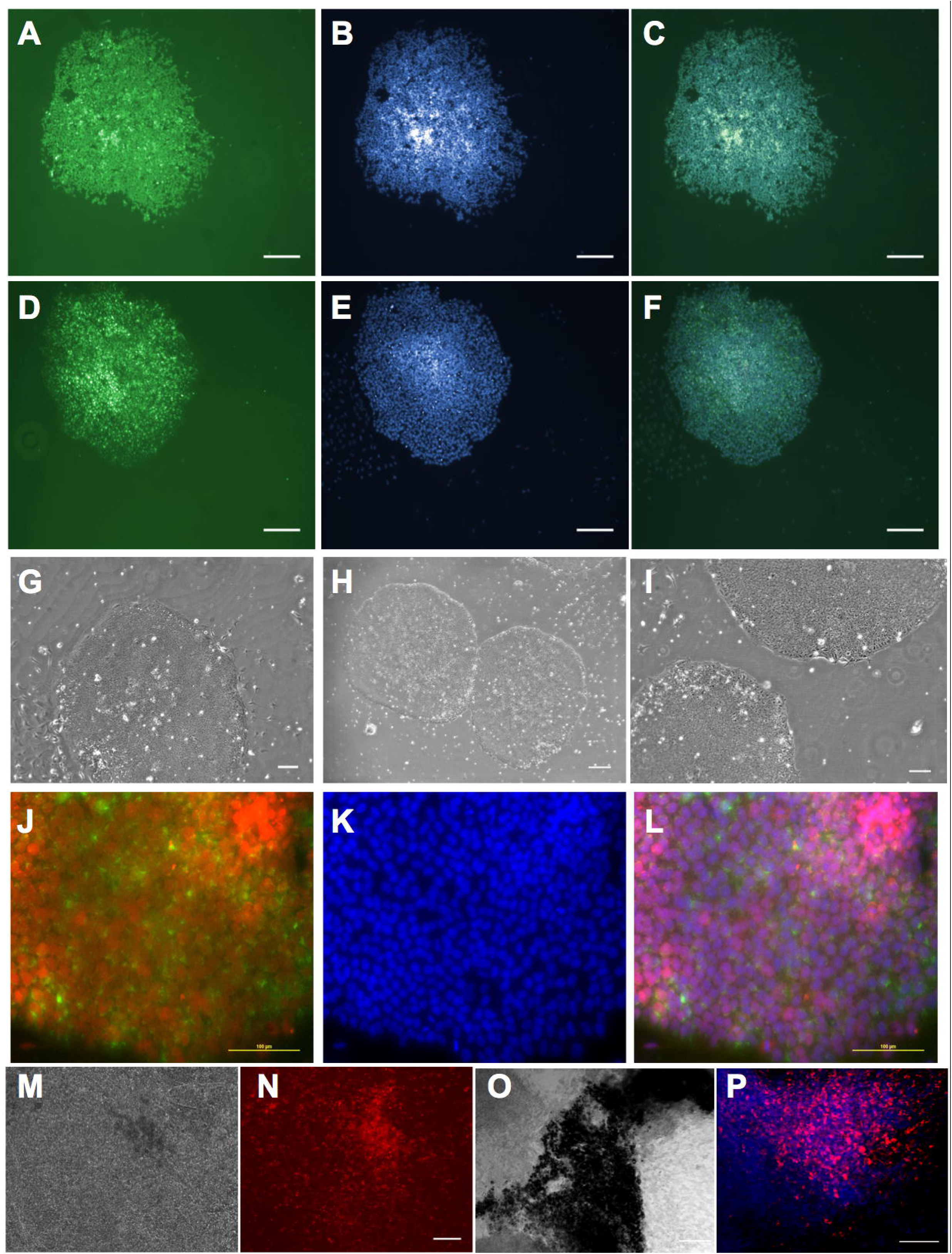
Stem cell maintenance and retinal differentiation using the automated platform. (A-F) Maintenance of cells post- TRA-1-selection. Representative images of colonies post-TRA-1-60 sorting, immunostained for OCT-4 (**A**) or TRA-1-60 (**D**) with DAPI counterstained (**B, E**) and merged (**C, F**). **(G-L) Passaging.** Representative bright field (**G-I)** and fluorescent (**J-L**) images of undifferentiated iPSCs after passage 1 (**G**), 3 (**H**) and 4 (**I**) and immunostained for OCT-4 (red) and TRA-1-60 (green) at passage 4 (**J**) with DAPI counterstained (**K**) and merged (**L**). (**M, N**) **RGC differentiation.** Representative bright field image (**M**) and corresponding fluorescence (**N**) of the reporter line BRN3B-mCherry A81-H7 following differentiation into RGCs at day 24. (**O, P**) **RPE cell differentiation.** Representative bright field (**O**) and fluorescent (**P**) images of H9 following differentiation to RPE cells at day 35 showing clear pigmentation and cobblestone morphology (**O**) and immunostained for the RPE marker PMEL with DAPI counterstain (**P**) characteristic of the RPE cells. Scale bars: **A-F**: 250 µm; **G, I-P**: 100 µm; **H**: 300 µm.

### Maintenance of PSCs

The maintenance and passaging templates allow for changing medium and passaging of iPSCs (Table 1). Maintenance was optimized for automation in serum-free and feeder-free conditions using vitronectin-coated plates in TeSR-E8 medium (Table 1). iPSCs were maintained for >5 passages on vitronectin-coated plates using E8 culture medium and passaged using ReleSR. Representative bright field images of colonies following successive passaging are shown in Fig. 2G-I. Immunocytochemistry confirmed that iPSCs remain pluripotent, as indicated by OCT-4 and TRA-1-60 expression, following successive passaging using this automated platform (Fig. 2J-L).

### Differentiation of PSCs into retinal cells on the automated platform

In order to assess the potential of this automated platform for long-term differentiation culture, we directed the differentiation of human PSCs towards two retinal lineages, RPE cells and RGCs, using protocols established within our group (Gill et al. 2016; Lidgerwood et al. 2016). We used the reporter line BRN3B-mCherry H7 (Sluch et al. 2015) for the RGC differentiation assay as this line fluoresces with expression of the specific RGC marker BRN3B, allowing for the screening of successful RGC differentiation. In order to make the RGC differentiation protocol of (Gill et al. 2016) suitable for automation, the protocol was slightly modified by replacing the initial embryoid body step with a monolayer differentiating culture. We then followed the protocol as previously described. As shown in Fig. 2M-N, we could clearly observe expression of mCherry in the differentiated culture, indicating successful RGC differentiation using our automated platform.

Next, we performed RPE differentiation using the automated platform. The differentiation of cells into RPE cells is evident by the characteristic morphology and pigmentation of RPE cells. We directed differentiation of human PSCs plated in feeder-free conditions into RPE cells using IGF1, DKK-1, noggin and bFGF as described in (Lidgerwood et al. 2016). Medium was changed every other day. Pigmented cells started appearing approximately four weeks later. We confirmed the polygonal geometry of the RPE cells and expression of the RPE marker PMEL (Fig. 2O, P). Together, these results provide proof-of-concept that automation can be utilised to facilitate stem cell maintenance and retinal differentiation. Importantly, no contamination was observed during the differentiation procedure, demonstrating the robustness of the automated platform for long-term sterile cell culture.

## Discussion

We describe the use of a modular platform to maintain, passage and differentiate human iPSCs. We adapted all protocols to automation using a feeder-free system for maintenance and differentiation. Some aspects of the work were performed offline, notably the reprogramming of cells and selection of successfully generated iPSCs. Further optimisation could allow these steps to be performed online, by integration within the modular platform, as done by others (Paull et al. 2015). We, however, decided not to integrate those in anticipation of potential changes in methods of reprogramming (in which nucleofection might become obsolete) or TRA-1-60 selection. The automated system allows for substantial customization of both equipment and cell handling parameters providing the flexibility needed for cell culture of various cell types. We did notice a few limitations of the automated platform. Despite a successful pipeline for the maintenance, passaging and differentiation of human PSCs, as well as sterility, improvements can be made to facilitate, streamline and reduce costs of the cell culture processes further. To reduce the risk of cross contamination, each line was cultivated within its own 6-well tissue culture plate. This format has the advantage of allowing for selections of cells for multiple applications, such as multiple long-term differentiation in various wells, or harvesting of samples for genomics, proteomics or lipidomics. On the other hand, this format could also imply that some wells within plates might stay empty, hence increasing unnecessary cost and waste. It would be very useful to also work with single-well tissue culture plates in addition to multi-well plates. However, those are currently not commercially available with the required height to be handled by the robotic arm of this platform. Such single-well tissue culture plates would be very beneficial as they could allow increasing a plating format as observed in flasks. Second, the conductive sterile filter tips are expensive. Disposable filtered tips are required, as fixed tips can easily carry contaminations and the sterility of the platform and in particular of the system liquid is paramount. There is also the risk of cross-contamination of samples by using fixed tips, thus we elected to solely use disposable tips for all liquid handling procedures. Also, maintenance of the platform and routine checks are essential but time consuming. Importantly, we have not had any single contamination using this platform, demonstrating the sterility of the system. Further, current templates for cell handling using the automated platform are relatively slow. Hence some adaptation had to be made to ensure cells were properly maintained, not left without medium for too long and placed back in the incubator in a timely manner. Future modification to the templates, such as reducing redundant movement of robotic arms, would reduce the time required for cell handling and increase the efficiency for the automated platform to process samples.

Finally, given the rapid pace of advances in the stem cell field, many assays and techniques for stem cell reprogramming, maintenance or differentiation can become rapidly obsolete and replaced with newer methods. Hence, a modular platform should allow for novel techniques to be included in the workflow to replace less useful methods. New protocols can be generated and adapted to the automated platform. However basic level of computing and training with the software are necessary. To fully realise the potential of the automated platform, the research group must be versatile and have a combined expertise in stem cell biology (reprogramming, cell maintenance and differentiation), as well as computing skills and engineering. To reduce these demands, workflow could be improved and made more generally useful by increasing the user friendliness of the interface between human and machine, thereby allowing modifications to the protocols to be made more easily.

The automated system we introduce here is novel and provides additional functions to the automated platform described by (Konagaya et al. 2015); it allows for the long-term maintenance and passaging of human iPSCs. In addition, we have also optimised maintenance of fibroblasts undergoing reprogramming as well as the directed differentiation to retinal lineages. This work can be placed in parallel to that of (Paull et al. 2015), who successfully used an automated platform for the generation, maintenance and differentiation of a large cohort of human PSCs. In their study, the authors used a remarkable 8 modular platform covering all aspects of cell handling. The system we describe here is not as complete but provides an economical alternative to the maintenance of PSCs and their differentiation.

In summary, we report the successful maintenance, passaging and differentiation of human PSCs, using an automated platform equipped with liquid handler and robotic arms, to handle cells in tissue culture plates. Despite some limitations, this platform shows excellent handling of sterility, and dramatically expands the potential of human PSC research. By increasing sample size and reducing variability, it allows for more defined parameters for future high-throughput analysis of the transcriptome and metabolome of progeny cells derived from patient iPSCs. This will be particularly important for iPSC modelling of complex genetic diseases, which will require large sample sizes to provide sufficient power for statistical analysis.

## Acknowledgements

We thank Marco Zalivani for the drawing of the CAD images of the automated platform.

## Author contributions

D.E.C., M.D., H.H.L., T.K., R.C.B.W.: concept and design, collection and/or assembly of data, data analysis and interpretation, manuscript writing, final approval of manuscript. F.L., G.E.L., A.C., D.H., S.S.H., K.P.G., E.D.S., L.K., L.C..: collection and/or assembly of data, data analysis and interpretation, editing and final approval of manuscript. V.M.S., X.C., D.J.Z.: generation of cells, editing and final approval final approval of manuscript. A.W.H., A.P.: concept and design, seeking of financial support, data analysis and interpretation, manuscript writing, final approval of manuscript.

## Conflict of interest/disclaimers

None

